# Mapping activity and functional organisation of the motor and visual pathways using ADC-fMRI in the human brain

**DOI:** 10.1101/2024.07.17.603726

**Authors:** Jasmine Nguyen-Duc, Ines de Riedmatten, Arthur P C Spencer, Jean-Baptiste Perot, Wiktor Olszowy, Ileana Jelescu

## Abstract

In contrast to blood-oxygenation-level-dependent (BOLD) functional MRI (fMRI), which relies on changes in blood flow and oxygenation levels to infer brain activity, diffusion fMRI (DfMRI) investigates brain dynamics by monitoring alterations in the Apparent Diffusion Coefficient (ADC) of water. These ADC changes may arise from fluctuations in neuronal morphology, providing a distinctive perspective on neural activity. The potential of ADC as an fMRI contrast (ADC-fMRI) lies in its capacity to reveal neural activity independently of neurovascular coupling, thus yielding complementary insights into brain function.

To demonstrate the specificity and value of ADC-fMRI, both ADC-and BOLD-fMRI data were collected at 3T in human subjects during visual stimulation and motor tasks. The first aim of this study was to identify an acquisition design for ADC that minimises BOLD contributions. By examining the timings in responses, we report that ADC 0/1 timeseries (acquired with b-values of 0 and 1 ms/*µm*^2^) exhibit residual vascular contamination while ADC 0.2/1 timeseries (with b-values of 0.2 and 1 ms/*µm*^2^) show minimal BOLD influence and higher sensitivity to neuromorphological coupling. Second, a General Linear Model was employed to identify activation clusters for ADC 0.2/1 and BOLD, from which average ADC and BOLD responses were calculated. The negative ADC response exhibited a significantly reduced delay relative to the task onset and offset as compared to BOLD. This early onset further supports the notion that ADC is sensitive to neuromorphological rather than neurovascular coupling. Remarkably, in the group-level analysis, positive BOLD activation clusters were detected in the visual and motor cortices, while the negative ADC clusters mainly highlighted pathways in white matter connected to the motor cortex. In the averaged individual level analysis, negative ADC activation clusters were also present in the visual cortex. This finding confirmed the reliability of negative ADC as an indicator of brain function, even in regions with lower vascularisation such as white matter. Finally, we established that ADC-fMRI timecourses yield the expected functional organisation of the visual system, including both gray and white matter regions of interest. Functional connectivity matrices were used to perform hierarchical clustering of brain regions, where ADC-fMRI successfully reproduced the expected structure of the dorsal and ventral visual pathways. This organisation was not replicated with the b=0.2 ms/*µm*^2^ diffusion-weighted time courses, which can be seen as a proxy for BOLD (via *T*_2_-weighting). These findings underscore the robustness of ADC time courses in functional MRI studies, offering complementary insights to BOLD-fMRI regarding brain function and connectivity patterns.

**Keypoints:**

1. The functional time course of the Apparent Diffusion Coefficient (ADC), specifically measured with alternating b-values of 0.2 and 1 ms/*µm*^2^ at 3T, appears to be minimally affected by BOLD contamination.
2. In the activity maps, the location of negative ADC clusters suggests neural activity in WM tracts that are connected to the motor cortex, which is not detected with positive BOLD.
3. Functional Connectivity analysis utilising ADC is better able to detect the organisation of the dorsal and ventral visual streams than diffusion- and *T*_2_-weighted time courses.

## Introduction

Functional Magnetic Resonance Imaging (fMRI) is a valuable tool in neuroscience research, providing insights into the function of both healthy and diseased brains. The primary contrast mechanism in fMRI is the blood oxygenation level-dependent (BOLD) signal [1, 2], which originates from localised variations in cerebral blood flow (CBF), volume and venous deoxyhemoglobin levels. These changes can manifest as either increases (positive BOLD) or decreases (negative BOLD) in the signal. Positive BOLD (pBOLD) is linked to neuronal activity through a process known as neurovascular coupling [3]. In task fMRI, negative BOLD (nBOLD) has been observed in sensory cortices not engaged by the stimulation paradigm, such as the deactivation of the auditory cortex during a visual task [4, 5]. However, it remains unclear whether nBOLD during a task reflects neural inhibition or results from a ‘blood-steal’ effect [6].

While the reliance of the BOLD signal on neurovascular coupling has proven informative, it introduces a significant constraint by not directly mirroring neuronal activity [7]. Consequently, this limitation brings about various challenges, including limited spatial and temporal specificity [8]. It is indeed typical to notice a delay between a stimulus and the BOLD response. Also, the BOLD signal does not emanate precisely from the location of neuronal firing, but rather from adjacent blood vessels [9, 10], which often leads to the overestimation of activated regions [11]. Furthermore, the study of neural activity in white matter (WM) encounters difficulties due to the scarcity of vasculature in these regions [12].

To address certain drawbacks associated with BOLD, diffusion MRI (dMRI) emerged as a promising alternative fMRI contrast. It leverages the hindrance and restriction of water molecules by micron-scale obstacles such as cell membranes, providing valuable insights into tissue cellular microstructure [13]. In the exploration of neural activity, several studies have reported a transient swelling and deformation of cells during neural firing [14]. This phenomenon could induce changes in the dMRI signal, which can be used as a diffusion functional MRI (DfMRI) contrast [15, 7, 8]. Neuronal swelling extends beyond the neuronal soma [16] to include axons [17, 18], specific dendritic regions [19], glial cells [20] and potentially even involves spines [21]. Prior studies suggest that these morphological alterations may result from cytoskeleton reorganisation [19] and/or from water influx through the cell membrane via an osmotic process [22, 23].“Neuromorphological coupling” is the term utilised to characterise the cellular swelling mechanisms observed in response to neural activity.

DfMRI’s sensitivity to neuromorphological coupling was supported by key observations such as the early onset of the DfMRI response (distinct from the delayed increase seen in BOLD responses) and a higher percent signal change with increasing b-values [7]. Such observations have been consistently found in various contexts, including in human [7], in rodent brain [24, 25, 23] and in cat brain [26]. Additionally, Tsurugizawa et al. [24, 27] demonstrated that nitroprusside, a drug suppressing blood reactivity to brain activity, significantly attenuated BOLD responses, while DfMRI responses showed minimal suppression as the neuronal response is preserved. This underscores the presence of information in DfMRI related to neuromorphological coupling, a facet not captured by BOLD signals.

However, there are indications in the literature that DfMRI contrasts using diffusion-weighted (DW) MRI timeseries may essentially be contaminated by changes in blood oxygenation and thus background field inhomogeneities, in the same way as BOLD. Notably, a study by Miller et al. [28] demonstrated significant modulations in the DW MRI signal during hypercapnia, a challenge that increases blood flow and volume but does not impact neural activity. This confound has been consistently observed in subsequent DfMRI studies utilising the DW signal time course [29, 24, 30]. It can be attributed to the intrinsic *T*_2_-weighted nature of the DW MRI signal, as dictated by the design of the spin-echo (SE) sequence. The alterations in tissue *T*_2_ due to variations in blood oxygenation levels during rest and activity form the fundamental mechanism driving the *T*_2_-BOLD contrast [7, 30, 28, 24, 10]. Consequently, a straightforward DW signal is significantly influenced by the same mechanism. However, there exist new approaches of ultra-high temporal resolution using line-scanning to separate early neuromorphological from later vascular effects. These approaches in the DW MRI signal have potentially singled out microstructural effects in the DfMRI contrast, that were detected before the delayed BOLD-like response [25, 27].

The dataset employed in our study originates in part from a prior investigation conducted by Olszowy et al. [10], where vascular contamination was minimised by a careful acquisition design. First, instead of directly using DW signals, the ADC at each timepoint is calculated from two DW volumes (with two different b-values), acquired in an interleaved fashion:

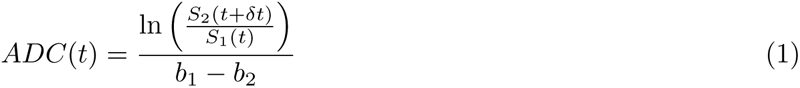

where *S*_1_ and *S*_2_ are the DW signals obtained with the b-values *b*_1_ and *b*_2_ respectively. The *T*_2_ varies on a much longer timescale than *δt*, therefore is assumed to be constant between *S*_1_(*t*) and *S*_2_(*t* + *δt*) [10] and essentially cancelled out in ADC. We term such an approach as ADC-fMRI from here on, to disentangle it from DfMRI which traditionally analyses the DW signal from a single b-value. To further remove vascular contamination, it has been reported that Twice Refocused Spin Echo (TRSE) sequence should be used instead of a Pulsed Gradient Spin Echo (PGSE) sequence, to minimise the cross-terms between the venous background field gradient (*G_s_*) and the diffusion gradient (*G_d_*) [31] :

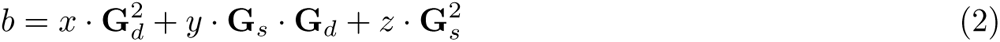

Previous studies have investigated neural activity through ADC time courses, recognising it as a compelling indicator due to its documented association with neural swelling during activity periods [15]. Consistent with DfMRI dynamics, ADC timeseries have exhibited an earlier variation following a stimulus compared to the BOLD response [27]. Le Bihan et al. reported a decrease in ADC timeseries with neural activity (negative ADC response) [15], while other studies have observed an increase (positive ADC response) [32, 33]. These discrepancies may reflect remaining BOLD contamination when using one b=0 measurement in the calculation of ADC. Indeed, studies reporting an increase in ADC typically calculated it with b_1_ as 0, whereas Le Bihan used a higher value of 0.2 ms/*µm*^2^. Including a b=0 measurement in the ADC estimation may yield response functions similar to BOLD effects, as observed in previous research [9], due to the direct contribution of blood water to the b=0 measurement but not to the one with a higher b-value (*b*_2_). Therefore, to suppress the contribution of fast-moving water molecules in blood to the signal, both b-values should be chosen equal to or above 0.2 ms/*µm*^2^ for the ADC calculation [10].

In this study, we extend the work of Olszowy et al. in validating ADC as a functional contrast mechanism, separated from BOLD and not influenced by neurovascular coupling. We separate negative ADC (nADC) from positive ADC (pADC) responses. We compare the spatial overlap in nADC vs pBOLD activation maps. Additionally, we examine the potential consistency of positive ADC (pADC) maps across subjects and whether they show spatial overlap with nBOLD. While numerous studies concur that, in task fMRI, nBOLD maps the deactivation of specific brain regions not involved in the task, such as the Default Mode Network (DMN) [34], to our knowledge, the use of pADC for this purpose remains unexplored. Similarly to Olszowy et al., we analyse ADC versus BOLD responses to simultaneous visual and motor stimuli, ensuring precautions to minimise BOLD contamination in the ADC time course. We also compare ADC responses derived from b-values of 0 and 1 ms/*µm*^2^ with those derived from b-values of 0.2 and 1 ms/*µm*^2^. We aim to identify potential signs of BOLD contamination absent in the “optimised” acquisition scheme.

Furthermore, we extend the analysis to functional connectivity (FC), particularly within the visual cortex. It is widely accepted that the visual cortex establishes connections with the broader brain through two main visual streams: the dorsal and ventral pathways. These pathways are recognised for their distinct functions crucial for ensuring precise visual perception [35, 36, 37]. Ungerleider and Mishkin [38] originally proposed this division based on anatomical findings in monkeys. Mishkin later concluded [37] that the dorsal stream processes information relative to spatial organisation (the “where”), while the ventral stream is primarily engaged in object identification (the “what”). At this stage of our investigation, we aim to assess whether ADC-fMRI could provide valuable insights into the FC within these visual pathways.

## Methods

### MRI experiments

A large part of the dataset employed in this study originated from a prior investigation conducted by Olszowy et al. [10]. Additional de novo datasets from 3 subjects were acquired for the current study using the TRSE ADC 0/1 protocol. Both studies obtained approval from the ethics committee of the canton of Vaud (CER-Vaud). A cohort of 18 participants was recruited in total, and all subjects provided written informed consent. Data were acquired on a Prisma 3T MRI scanner (Siemens Healthineers, Erlangen, Germany) equipped with 80 mT/m gradients and fitted with a 64-channel receiver head coil.

The study implemented 3 scanning protocols:

1. SE-EPI for T2-BOLD contrast.
2. DW-TRSE-EPI with b-values 0.2 and 1 ms/*µm*^2^ (TRSE ADC 0.2/1).
3. DW-TRSE-EPI with b-values 0 and 1 ms/*µm*^2^ (TRSE ADC 0/1).

Sequences were obtained from the Center for Magnetic Resonance Research at the University of Minnesota (https://www.cmrr.umn.edu/multiband/). Details on the acquisition parameters can be found in Table 1. All scans employed GRAPPA=2 and Multiband=2 acceleration. The two b-values were acquired alternately within the same functional run, producing an ADC map with half the temporal resolution (TR) of SE-BOLD. The TR for SE-BOLD was one volume per 1.035 seconds for all subjects. For DWI-TRSE-EPI, the TR was one volume per 1.035 seconds when the alternating b-values were 0.2 and 1 ms/*µm*^2^, and was one volume per second when the alternating b-values were 0 and 1 ms/*µm*^2^. The short TR of the sequences could only yield partial brain coverage. The imaging slab covered both the visual and motor cortices, with outermost slices discarded due to signal hyperintensity and to possible inflow contributions. To facilitate later susceptibility distortion correction using topup [39], additional EPI volumes were acquired with an opposite phase encoding direction. Additionally, a whole-brain 1 mm isotropic T1-weighted anatomical image was obtained using an MPRAGE sequence.

During the task, subjects viewed a flashing checkerboard (8 Hz) while concurrently finger-tapping with both hands for 12s, following 18s of rest. Each block was repeated 20 times per run, resulting in a total acquisition time of 10 minutes per subject (Figure 1). This paradigm was implemented using PsychoPy [40].

**Table 1:** Acquisition parameters for the 3 fMRI protocols.

|  | SE-BOLD | DW-TRSE-EPI 0/1, TRSE | DW-TRSE-EPI 0.2/1, TRSE |
| --- | --- | --- | --- |
| # Datasets | 4 | 4 | 8 |
| TE (ms) | 73 | 72 | 72 |
| TR (s) | 1.035 | 1 | 1/1.035 |
| Matrix | 94x94 | 116x116 | 94x94 |
| Resolution (mm<sup>2</sup>) | 2.5x2.5 | 2x2 | 2.5x2.5 |
| Slices | 16 |  |  |
| Thickness (mm) | 2.5 | 2 | 2.5 |
| Total volumes | 600 |  |  |

### Data Processing

The data processing pipeline was developed in Python, following a methodology similar to the preprocessing described in [10], which was implemented in Matlab. The pipeline comprised outlier detection, MP-PCA denoising, Gibbs unringing, top-up correction, motion correction, and the generation of ADC maps.

#### Detection of volume outliers

Signal dropouts, manifested as outliers in the timeseries, were automatically detected and replaced with linearly interpolated signals. A volume was flagged as an outlier if its linearly detrended average signal across the entire brain deviated by more than 1% from the median across all volumes. Typically, the proportion of outliers remained below 5%. However, if this proportion exceeded 20%, the entire dataset was omitted from subsequent analyses. For the ADC-fMRI datasets, outlier detection was performed separately for volume series corresponding to the two b-values.

#### MP-PCA Denoising, Gibbs Unringing, and Top-up

The denoising process employed the Marchenko-Pastur Principal Component Analysis (MP-PCA) [41] approach within MRtrix3 [42]. For ADC-fMRI datasets, an initial split into two distinct arrays was first necessary, one for each of the two alternating b-values. The denoising procedure was then applied to these arrays separately [43], to avoid subtle changes in amplitude being removed along with the thermal noise. BOLD-fMRI time courses were also denoised using MP-PCA [44]. Additionally, corrections were implemented to address the Gibbs ringing artifact [45] and susceptibility distortions [39] in BOLD and ADC acquisitions, with the latter being resolved using FSL’s [46] topup.

#### Motion Correction

Motion correction was conducted using the ANTs (Advanced Normalisation Tools) library [47]. For ADC-fMRI datasets, the input was divided into two arrays representing different b-values. The mean of each array served as a reference for motion correction. The mean *b*_2_ image was registered to the mean *b*_1_ image. Subsequently, motion correction was independently applied to each array using the “antsMotionCorr” function with rigid transformation. The resulting motion correction transformation was then concatenated with the *b*_2_ to *b*_1_ transformation to align the second array with the first in space. This produced the motion-corrected ADC-fMRI data. Motion correction was similarly applied to BOLD-fMRI data, without the prior split into distinct arrays.

#### Creation of ADC maps

From the preprocessed ADC-fMRI datasets, ADC time courses were calculated using equation 1. ADC values exceeding 4 *µm*^2^*/ms* or falling below 0 were treated as outliers and were substituted with 0. These outliers were found to be exclusively outside the brain for all subjects.

#### Atlas registration

To enable group-level analysis and atlas-based region-of-interest (ROI) segmentation, functional data were registered from native space to MNI atlas space using ANTs. Following brain extraction (using FreeSurfer’s synthstrip method [48]), the subject’s anatomical image underwent registration to the Montreal Neurological Institute (MNI) template at a 1 mm isotropic resolution using symmetric normalisation (non-linear registration). Subsequently, the mean functional image was aligned with the subject’s anatomical image using a rigid transformation. In the case of ADC-fMRI data, the mean functional image was computed from the lower b-value volumes. The rigid and non-linear transformations were applied to bring individual functional maps into atlas space, or atlas ROIs into native space.

**Figure 1:**
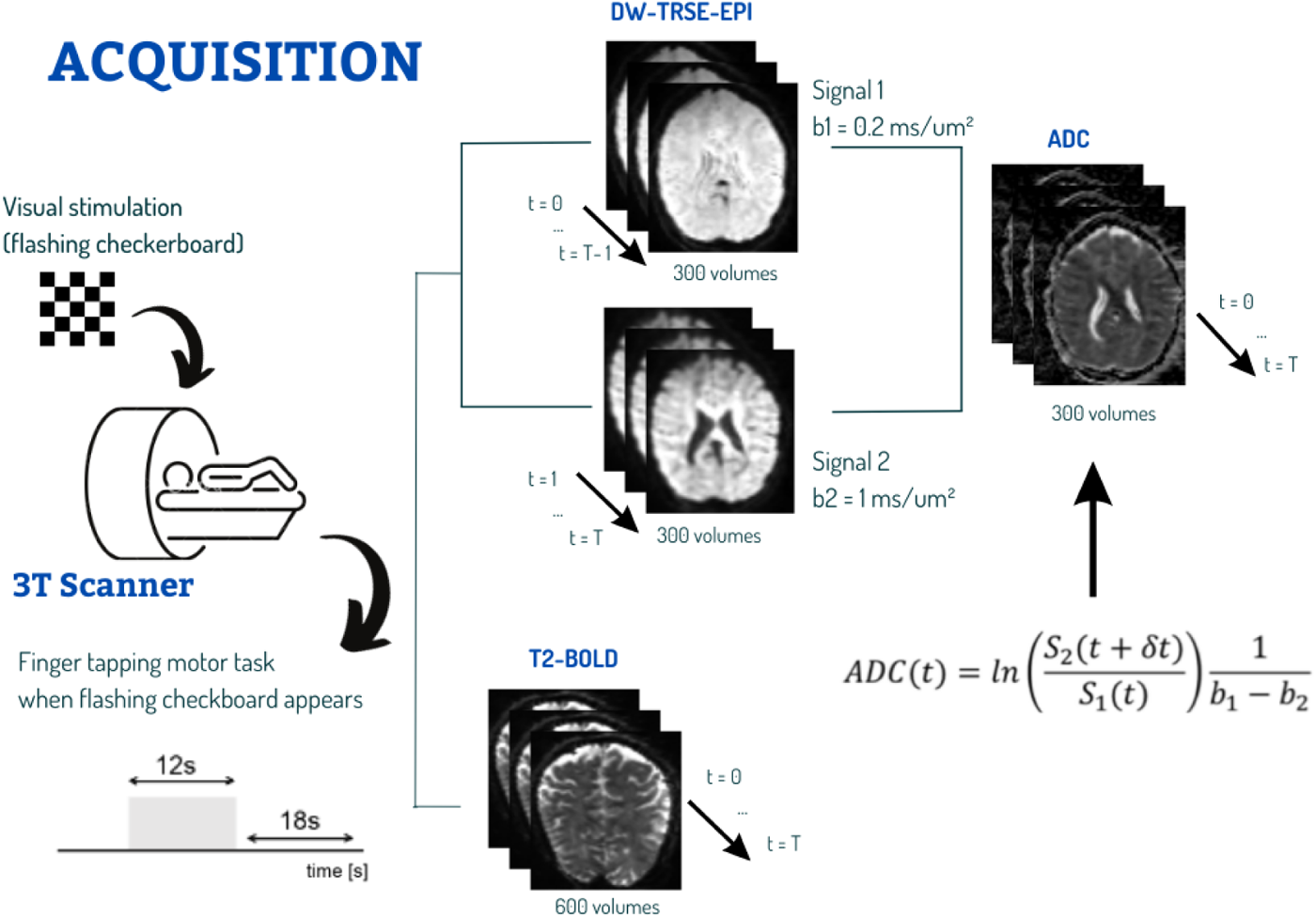
Subjects underwent imaging sessions on a 3T MRI scanner. The fMRI data acquisition employed distinct imaging modalities. The DW MRI timeseries with interleaved b-values, yielded ADC time courses with 300 volumes. The SE-EPI timeseries yielded BOLD fMRI time courses with 600 volumes. During the imaging sessions, a visual and motor task utilising a flashing checkerboard with a frequency of 8 Hz, with simultaneous finger-tapping, was repeated in 20 blocks.

### ROI-averaged ADC 0/1 vs 0.2/1 timeseries

Prior to any GLM analysis, we examined ADC time courses averaged across ROIs expected to be involved in the visual and motor tasks: V1, V2, BA4a, BA4p, BA1 and BA2, left and right, as defined by the Juelich atlas [49] in native space. The objective was to compare levels of vascular contribution in the ADC 0/1 vs 0.2/1 timeseries. The time courses were normalised with respect to a baseline, defined as the 7 seconds preceding the task, and then averaged across ROI voxels, epochs and subjects. Subsequently, for each of these averaged time courses, a Linear Regression was calculated between the time course and both the boxcar function and the canonical HRF convolved with the stimulus. The resulting p-values were adjusted for multiple comparisons using Bonferroni correction. Only significant correlations (*p <* 0.05) were reported.

### ADC 0.2/1 vs BOLD GLM analysis

GLM analyses were conducted using FEAT [50] in FSL [51, 52] on ADC 0.2/1 and BOLD data in native space. We used a boxcar response function to avoid assumptions about response shapes, which are unknown for ADC-fMRI. For BOLD response analysis, the HRF is typically used. However, using the same analysis pipeline for both BOLD and ADC facilitated a direct comparison. Subject-level activation maps were computed using cluster correction (*|z| >* 2.3, *p <* 0.05). Significant voxels were categorised as positively correlated with the task (*z >* 2.3) or negatively correlated (*z < −*2.3). The time courses in positively and negatively correlated voxels of the visual and motor cortices were normalised to the baseline within each epoch separately. Normalised timeseries were then averaged across epochs and subjects to yield averaged BOLD and ADC-fMRI responses to the task. The visual and motor cortices were defined in subject space using the 1.5 mm isotropic Neuromorphometrics atlas, as in [10]. The visual cortex was the sum of the calcarine cortex, occipital pole, superior occipital gyrus, inferior occipital gyrus, cuneus, lingual gyrus, middle occipital gyrus and occipital fusiform gyrus. The motor cortex was the sum of the precentral gyrus medial segment, precentral gyrus and supplementary motor cortex. All the regions were considered bilaterally.

To further investigate the differences in responses between BOLD and ADC, a Mann-Whitney U Test [53] was employed with a p-value threshold of 0.05. This test assessed whether the delay from task onset to signal change, and the delay from task offset to return to baseline, were significantly greater for BOLD than for ADC. The timing of the signal change was defined as the time after task onset that the signal exceeded the mean for the epoch (or dropped below for negative responses). Similarly, the timing of the return to baseline was defined as the time after task offset that the signal dropped below the mean value for the epoch (or above for negative responses). The timeseries considered were averaged across epoch and per voxel for each subject to obtain a smooth timeseries to easily compute the timings mentioned. All significant voxels (*|z| >* 2.3*, p <* 0.05) were taken into account, regardless of the brain region they belonged to.

To assess spatial activation maps, individual cluster-corrected z-score maps were transformed to MNI space and averaged across subjects. Additionally, a group-level analysis was performed using FMRIB’s Local Analysis of Mixed Effects (FLAME) [54] with stages 1 and 2, with a cluster correction (*|z| >* 1.5, *p <* 0.05) to control for false positives. Group-level activation maps were computed for both BOLD and ADC 0.2/1, again categorising clusters as positively correlated (*z >* 1.5) or negatively correlated (*z < −*1.5) with the task.

### ADC and DW Functional Connectivity analysis

For this analysis, FC matrices first were computed for each subject with the ADC 0.2/1 timeseries and the DW timeseries (b = 0.2 ms/*µm*^2^). FC between ROIs was calculated using partial Pearson correlations, with the average cerebrospinal fluid (CSF) time course (defined by the ventricle ROIs of the Neuromorphometrics atlas) serving as a covariate. An averaged FC matrix was obtained for each contrast. The DW timeseries was used in this analysis instead of the BOLD timeseries to ensure that the time courses originate from the same functional run. This approach, where the b = 0.2 ms/*µm*^2^) measurements are a subset of those underlying the ADC 0.2/1 time course, helps to minimise intra-subject variability in FC. The aim was to compare FC analysis with the ADC, which is expected to be minimally influenced by vascular components, against a time course known to be affected by T2 weighting (and thus BOLD contrast), which is the DW timeseries.

The ROIs were defined using the Juelich and John Hopkins University (JHU) [55] atlases of GM and WM, respectively. All WM ROIs in the Juelich atlas were discarded, except for the Optic Radiation, which is not present in the JHU WM atlas. Supplementary Table 1 shows the ROIs that are taken into account in this work. They were transformed to subject space using ANTs, so that the temporal dynamics of the signals within each ROI could be averaged in subject space. If at least 10% of the ROI was covered by the slab for all subjects, the ROI was considered for the analysis.

To organise the ROIs in an informative manner, a tree structure was created for each FC matrix using Python Seaborn’s implementation of the UPGMA algorithm for a hierarchical clustering [56]. This algorithm facilitated the hierarchical arrangement of ROIs in a tree-like structure. The goal was to uncover an organisational scheme that illustrates how the brain’s ROIs can be arranged based on the similarity of their connections to the rest of the brain. Additionally, we aimed to identify whether the FC patterns associated with the dorsal and ventral visual streams could be discerned. For this, the FC matrix was adjusted to reflect the connectivity of each ROI (from the left and right hemispheres) to their corresponding ipsilateral and contralateral ROIs. ROIs that were not bilateral (such as the forceps major) were discarded.

To visualise connections pertinent to the two-stream hypothesis, a graph was constructed from the FC matrix obtained (before the latter adjustments) with the ADC data. Pairs of ROIs with correlation values exceeding 0.4 were considered and saved as edges of this graph. Edges that did not involve any ROIs in the visual cortex, as defined by the Juelich atlas (including V1, V2, V3v, V4, and V5), were excluded.

## Results

### Comparison between ADC 0/1 vs 0.2/1 ROI-averaged responses

Figure 2 presents the averaged responses for ADC 0/1 and ADC 0.2/1 across voxels, epochs, and subjects within 12 ROIs where neural activity was anticipated. When including all voxels in a given anatomical ROI, changes in the ADC 0.2/1 level during the task were mostly undetectable by eye. Conversely, the ADC 0/1 response in the visual cortex ROIs (V1 and V2) exhibited a distinct pattern: the signal decreased at task onset, then increased after approximately 5 seconds, and eventually returned to baseline after a delay of 8 seconds following the task offset. The amplitude of this increase reached 1%. Figure 3 illustrates the slopes from the Linear Regression (adjusted for multiple comparisons) between each averaged time course (from Figure 2) and two response functions: the boxcar function and the canonical HRF convolved with the task. In 8/12 ROIs, the averaged ADC 0/1 time course was significantly positively correlated with the HRF, with slope values that varied between 0.0013 and 0.011. No significant correlation was found between ADC 0/1 time course and the boxcar function. The averaged ADC 0.2/1 time course showed significant negative correlations with the boxcar function in 4/12 ROIs, and a significant negative correlation with the HRF in one ROI. The negative slopes varied from -0.0013 to -0.003.

**Figure 2:**
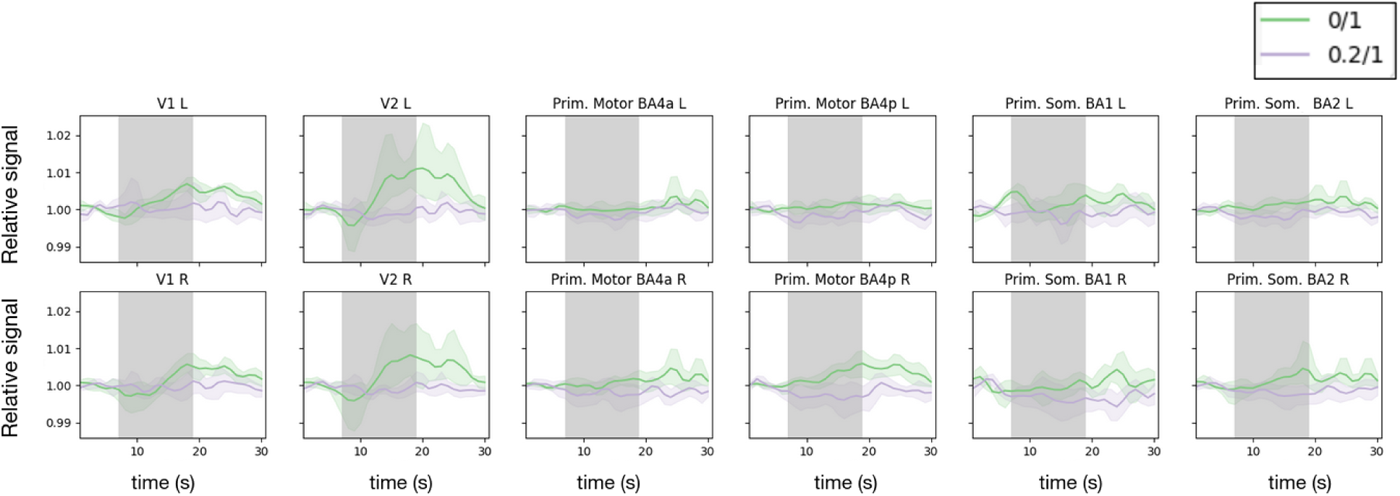
Averaged ADC responses with b-values of 0/1 ms/*µm*^2^ (green) and 0.2/1 ms/*µm*^2^ (purple) are displayed for specific ROIs in the visual, motor, and somatosensory cortices, where neural activity is anticipated during the task. The shaded areas around each curve represent the 95 % confidence band, illustrating the variability between subjects (8 for ADC 0.2/1, 4 for ADC 0/1).

### ADC 0.2/1 vs BOLD response functions

The average positive and negative responses originating from significant voxels obtained from the subject-level GLM analyses are shown for the visual and motor cortices in Figure 4. The amplitude of the pBOLD response exceeded that of the nBOLD response (2% versus 1% in the visual cortex and 1% versus 0.5% in the motor cortex), with a greater percentage of voxels contributing to the positive response compared to the negative response (approximately 80% versus 20%).

Regarding ADC 0.2/1, both positive and negative responses displayed approximately 4% amplitude in the visual cortex and 2% in the motor cortex, which is remarkably larger than BOLD. However, the 95% confidence band, as shown by the shaded area, implies higher variability across subjects in the ADC timeseries compared to the BOLD timeseries. A greater number of voxels contributed to the negative response than to the positive response (70% versus 30% in the visual cortex and 90% versus 10% in the motor cortex). Comparing counts between functional contrasts, ADC-fMRI yielded five to ten times fewer significant voxels than BOLD (pBOLD vs nADC, and nBOLD vs pADC).

Despite being analysed using a GLM with a boxcar function, the BOLD average response exhibited the expected hemodynamic delay whereas ADC did not. Figure 5 shows that this delay was significantly shorter in nADC compared to pBOLD for both the delay from task onset to signal rise/fall (*p* = 0.03) and the delay from task offset to return to baseline (*p* = 0.01). The average delays were around 5 seconds for pBOLD and 0 to 1 second for ADC. nBOLD displayed a delay primarily in return to baseline after task offset.

**Figure 3:**
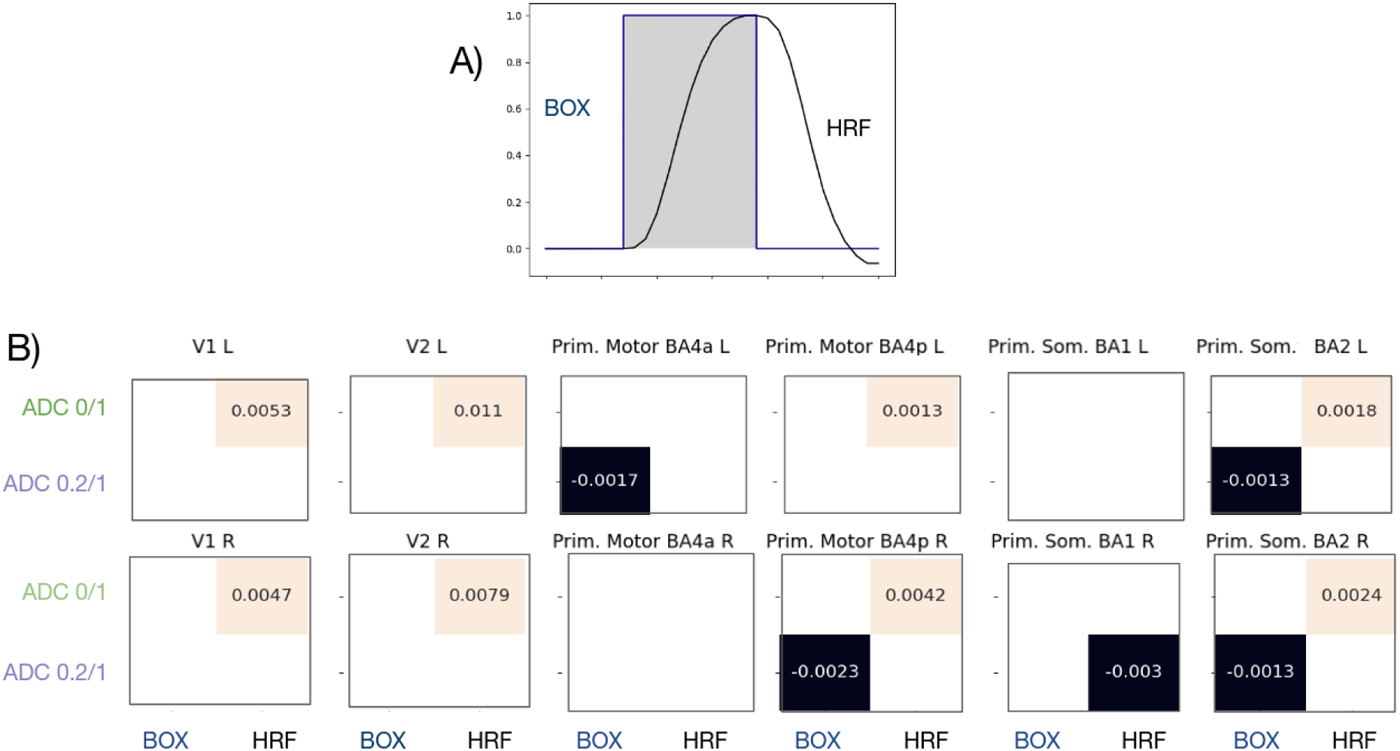
A) The boxcar function (BOX) and the canonical HRF convolved with the task. B) Pearson correlations were computed between the averaged ADC timeseries (0/1 and 0.2/1) with the BOX function and convolved HRF. The p-values were corrected for multiple tests using the Bonferroni correction, and non-significant results (*p >* 0.05) were omitted.

### Averaged cluster-corrected z-score activation maps

Figure 6 presents the cluster-corrected z-score maps averaged across individuals. In Figure 6A, the pBOLD effectively depicted neural activity in the visual and motor cortex, as expected. The resulting clusters were large, with high averaged z-scores, ensuring spatial overlap between significant clusters across different subjects. As Figure 6B illustrates, clusters exhibiting nADC were notably smaller, predominantly unilateral (in the right hemisphere) and displayed limited overlap across subjects, as shown by the lower averaged z-score. They were primarily located in the visual cortex, as expected due to the visual stimulus. Interestingly, nADC clusters were found outside of the visual and motor cortices, in the WM connecting the motor cortex to the corticospinal tract (CST) and the inferior parietal lobe to the superior parietal lobe.

Clusters of negative BOLD response were found within the visual cortex, in distinct locations from those of pBOLD. These nBOLD clusters were situated above the calcarine fissure and closer to the parietal cortex. On the other hand, clear pADC clusters were challenging to discern, appearing very small and scattered.

### Activation maps from group level analysis

Figure 7 shows the cluster-corrected group level analysis. These maps show a more conservative analysis, with fewer significant clusters than what was found in Figure 6. Figure 7A presents the pBOLD and nADC, supposedly corresponding to excitatory response. The pBOLD clusters were in identical regions, as previously shown in Figure 6, but were spatially more limited. The significant nADC voxels that were present in the visual cortex were no longer present. However, a pathway found in WM that was connected to the motor cortex remained. The voxels showing nBOLD clusters (in Figure 7B, associated with inhibitory response) were found only in the right angular gyrus, which is a region that is part of the DMN. No pADC voxel cluster survived the group-level analysis.

**Figure 4:**
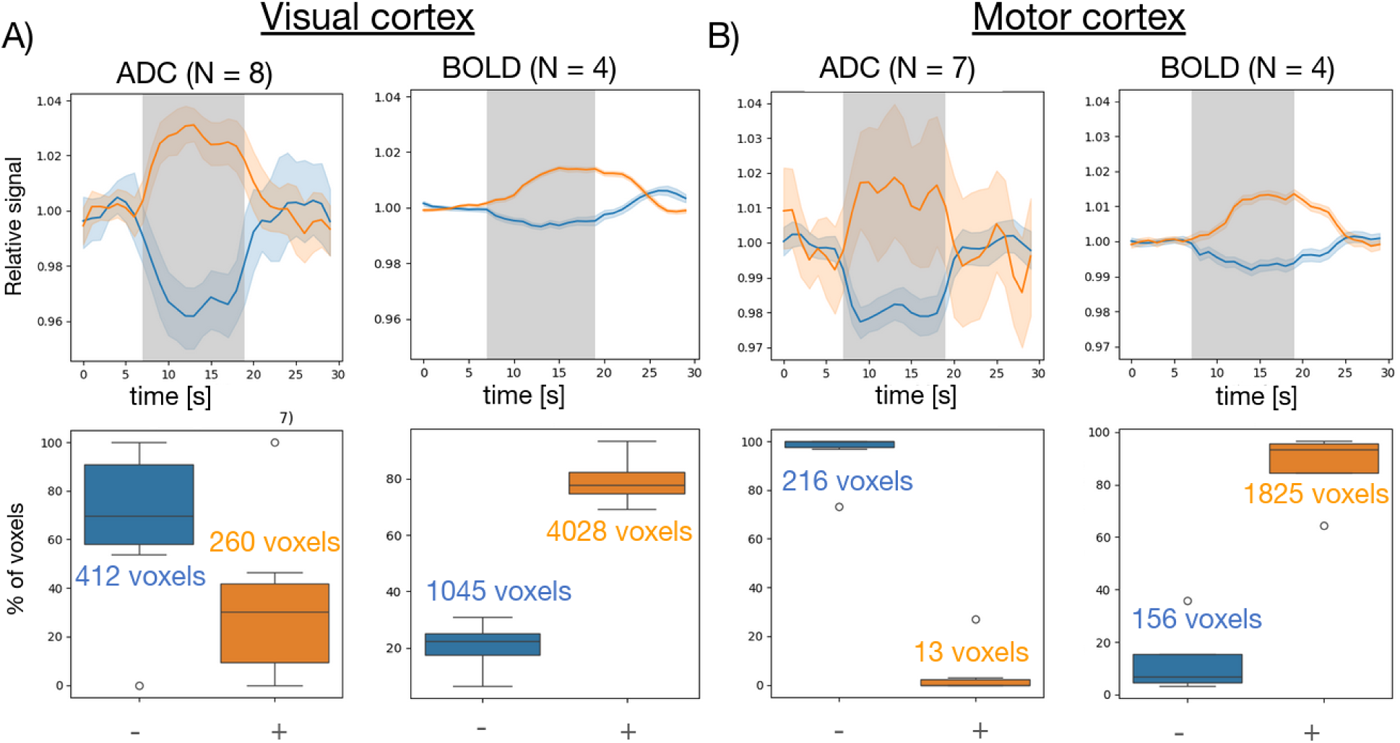
The averaged ADC and BOLD responses stem from voxels exhibiting significant correlation with the task, identified through a GLM analysis with cluster correction (*|z| >* 2.3, *p <* 0.05) conducted on each subject. timeseries are averaged across epochs and subjects, with the orange curve representing the average timeseries positively correlated with the task and the blue curve depicting the average of negatively correlated timeseries. Shaded areas around each curve indicate the 95% confidence band, demonstrating variability between subjects. The box plot below illustrates the percentage of voxels contributing to the positive and negative curves, with the deviation indicating variability among subjects. These results are provided for both the visual (A) and motor (B) cortices.

### ADC-fMRI Functional Connectivity

Upon examining FC, the hierarchichal clustering presented in Figure 8 and in Supplementary Figure 1 illustrated hierarchical tree structures generated using the UPGMA algorithm. These trees organised ROIs based on the similarities of their connections to other ROIs within the brain slab (which were relatively ipsilateral or contralateral). Figure 8 shows the results obtained with ADC-fMRI, whereas Supplementary Figure 1 shows the results when DW timeseries (b = 0.2 ms/*µm*^2^) are utilised instead. Areas involved in similar functions to each other are expected to belong to the same branch of the tree. For example, the trees derived from both the ADC and DW data distinguished ROIs of the visual cortex in a separate branch from the rest of the ROIs. In Figure 8, the visual cortex branch was further divided into distinct branches containing the V3v/V4 areas, the V5 areas, the V1/V2 areas, and the Optic Radiation (Optic Rad)/Inferior fronto-occipital fasciculus (IFO)/Inferior longitudinal fasciculus (ILF), which are the white matter tracts involved in the visual system. Regarding the “non-visual” branch (on the right side of the tree), there was indeed a clear organisation of ROIs into different sub-branches based on their functions, such as the Inferior Parietal lobe (IPL), the Anterior intra-parietal sulcus (AIPS) along with remaining ROIs of the IPL, the Somatosensory/Motor cortices and finally the Superior longitudinal fasciculus (SLF) along with the Cingulum (see names of ROIs and their shorter version in Supplementary Table 1). In Supplementary Figure 1, given the T2-weighting of the b = 0.2 ms/*µm*^2^ DW-fMRI time course, identifying functional clusters within each branch of the tree was more challenging. The organisation of ROIs appeared less systematic and more arbitrary, with ROIs that were expected to be grouped together often found in separate branches. For instance, the AIPS ROIs were not grouped together in a branch as they were in the ADC FC hierarchical clustering. Additionally, within the visual tree, V2 R is found in the same branch as the WM tracts within the visual cortex, rather than being positioned near V2 L and the V1 ROIs, as would be expected. This lack of clear clustering contrasted with the more logical arrangement observed in Figure 8, making it difficult to discern any coherent pattern or functional relationship among the ROIs.

**Figure 5:**
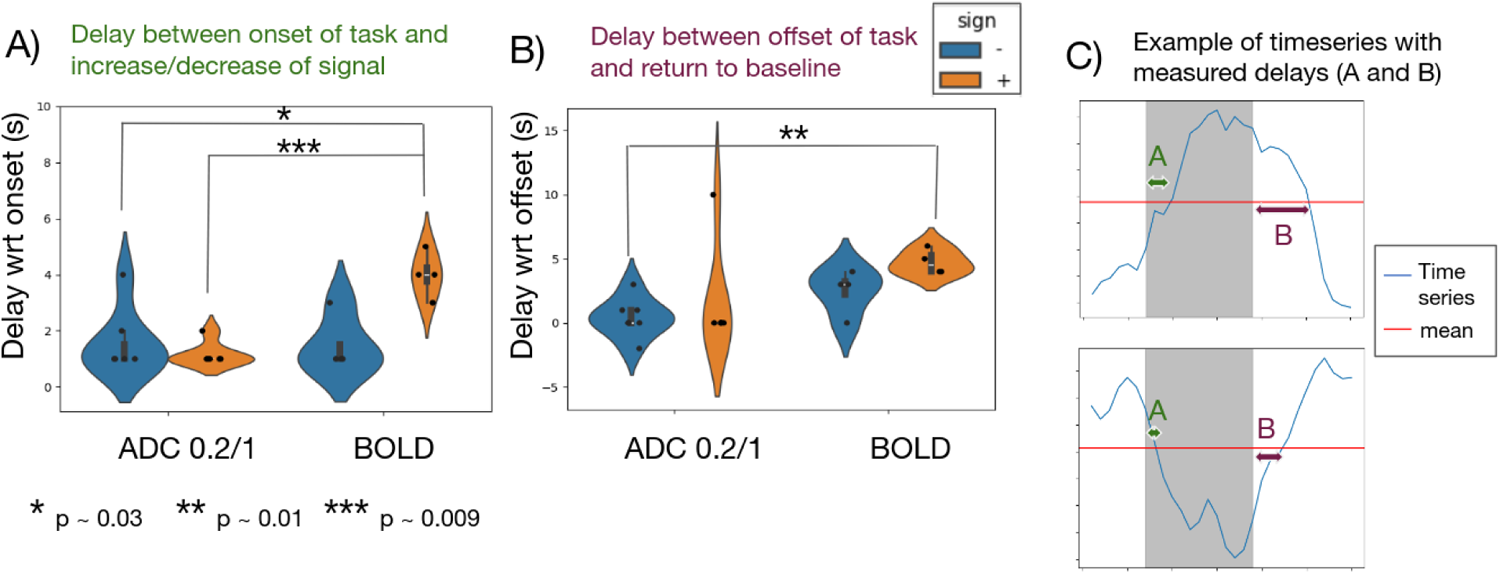
The violin plots shows the delays relative to the task for ADC 0.2/1 and BOLD. A) The delay was calculated from task onset to the signal change (the first time the signal surpasses or drops below its mean value during the task period). B) The delay was measured from task offset to the return to baseline (the second time the signal surpasses or drops below its mean during the following rest period). Statistical analysis was conducted using Mann-Whitney U Tests in both A and B. C) Example of timeseries (averaged across voxels and epochs for a random subject), illustrating how the two types of delays (denoted by A and B) are measured. Delay A corresponds to the delay depicted in (A), and delay B corresponds to the delay depicted in (B).

In the concluding stage of our analysis, we established an ADC-fMRI derived graph illustrated in Figure 9 with the specific aim of highlighting the visual ROIs graph (V1, V2, V3v, V4 and V5) obtained from Pearson partial correlations exceeding a threshold of 0.4. The analysis confirmed that ROIs corresponding to WM tracts present in the “visual branch” of the tree in Figure 8 exhibited considerable connections to these visual cortical ROIs. Additionally, the Forceps Major displayed strong connections to V1 and V2, consistent with its location in the occipital cortex and its role as a bridge between the hemispheres. Furthermore, two IPL ROIs were identified: the IPL Pgp, which was connected to most visual ROIs, and the IPL Pga (present in the angular gyrus), which was connected only to V5. Remarkably, a possible transmission of information can be observed in both hemispheres: starting from V1, the information flows to V2, then progresses to V3v, subsequently connecting to V4, and finally reaching V5. The overall graph demonstrated a highly symmetrical shape, with connections in the left hemisphere mirrored by those in the right hemisphere.

**Figure 6:**
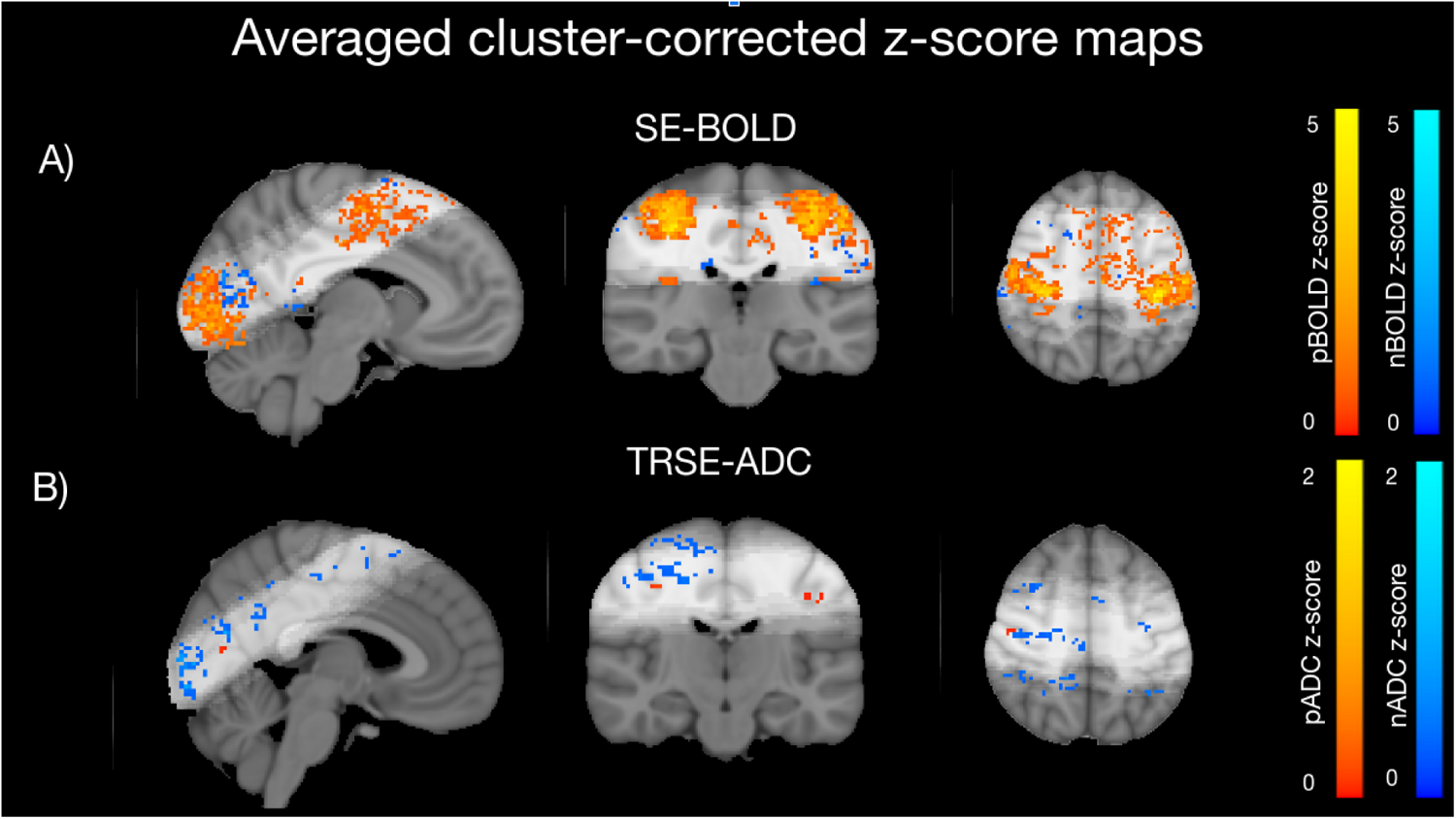
Voxels demonstrating significant correlation after cluster correction (*|z| >* 2.3, *p <* 0.05) with the task were identified to depict neural activity using both BOLD (A) and TRSE ADC 0.2/1 (B). The visualisation presents voxels with timeseries that negatively correlate with the task in blue (nADC, nBOLD) and those that correlate positively in red (pADC, pBOLD). The scales indicate the average z-score across all subjects. The light shaded area in the brains show the slab from which data was acquired.

## Discussion

### Assessing Vascular Contamination in ADC Responses Based on DfMRI Protocols

One of the objectives of this investigation was to assess the potential of ADC as an independent functional MRI contrast, unaffected by vascular contributions. Our hypothesis was centred on previous evidence that ADC, specifically with b-values of 0.2 and 1 ms/*µm*^2^, would effectively capture neural activity through neuromorphological coupling, exhibiting minimal contamination from vascular effects. On the other hand, in ADC 0/1 it was previously hypothesised that water molecules in blood contribute to the response by influencing the b = 0 signal but not the b = 1 ms/*µm*^2^ signal, as the latter is not sensitive to fast flow. In this scenario, ADC sensitivity extends beyond the swelling of neurons and glial cells to include the expansion of arteries, veins, and capillaries during increased blood flow. This theory is supported by a study by Le Bihan et al. [7], which notes that DW MRI timeseries with *b*_1_ = 0.2 exhibit lower amplitude than those with *b*_1_ = 0. To test our hypothesis, we compared the two different ADC responses in multiple ROIs where neural activity is expected.

The ADC 0/1 response in the visual ROIs (V1 and V2) exhibited a distinct pattern: an initial decrease in ADC was quickly followed by an increase 5 seconds into the task, with a return to baseline occurring 8 seconds after the task ended. In [27], where the ADC time course calculated can be compared to our ADC 0/1 time course, a similar pattern in the signal was observed. The rapid onset of the initial dip was interpreted as evidence of the ADC time course’s sensitivity to neuromorphological coupling. The subsequent signal increase was attributed to a BOLD effect, possibly convolved with a ‘glio-morphological’ effect [27]. This increase made the ADC 0/1 response resemble the convolved HRF, leading to strong correlations between the time course and the convolved HRF response in most of the 12 ROIs (8/12).

In the averaged ADC 0.2/1 response, this delayed increase pattern mirroring the BOLD response was not observed. We expected to find a decrease in the response with neural activity during the task. This decrease was difficult to discern, likely due to the combined effects of limited ADC-fMRI sensitivity and the averaging of all voxels within each ROI, including sub-regions that may not respond to the task. Despite this, significant negative correlations were found between ADC 0.2/1 and the boxcar function in 4/12 ROIs. This indicates that, although the decrease is small in amplitude, it is meaningful and not attributable to randomness. Since the time course was significantly negatively correlated with the boxcar function more frequently than with the convolved HRF (4/12 versus 1/12), it is very likely that ADC 0.2/1 did not exhibit vascular contamination. This implies that it is driven by early neuromorphological coupling, without the influence of the later BOLD effect, which was observed in ADC 0/1.

**Figure 7:**
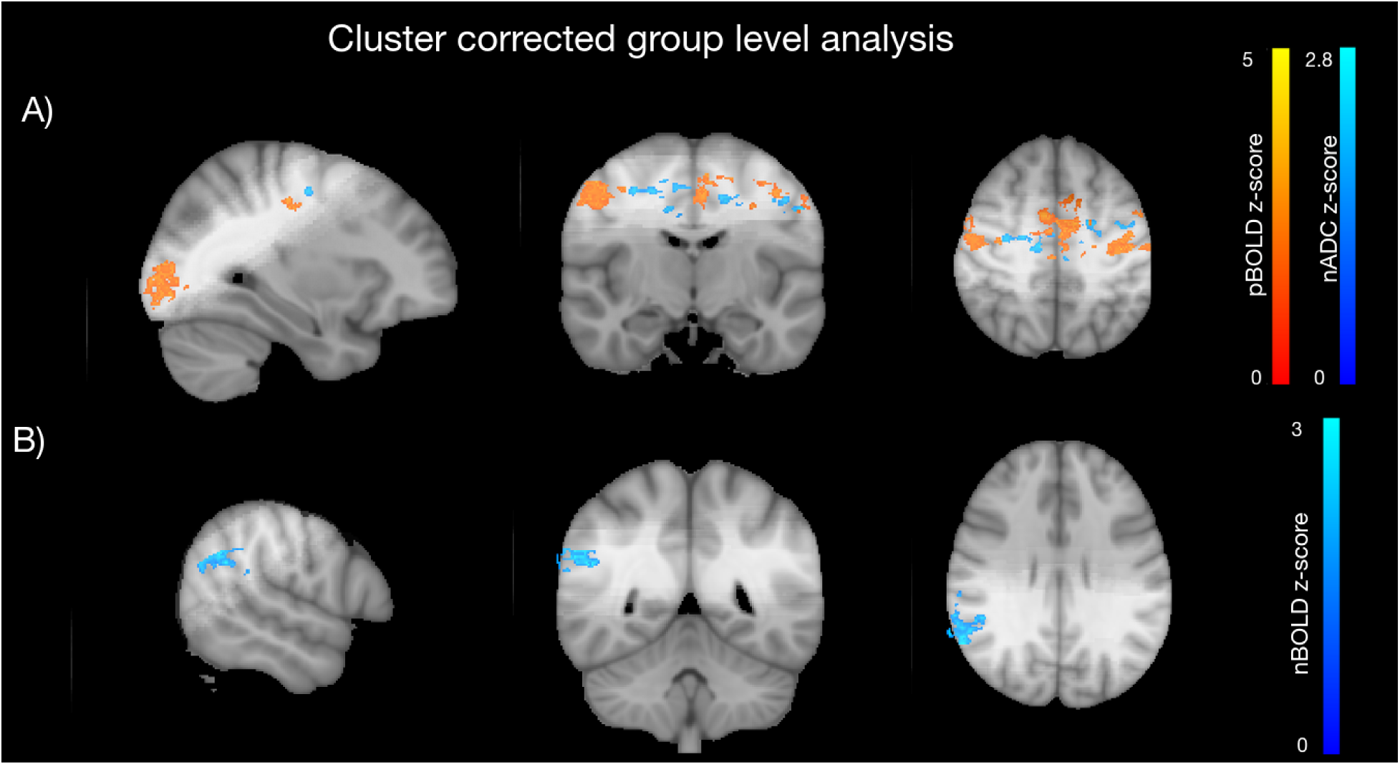
Group-level analysis was conducted using FLAME (stages 1 and 2) for both BOLD and TRSE ADC 0.2/1 data. Cluster correction was applied using *|z| >* 1.5 and *p <* 0.05. Neural activity is depicted with pBOLD (orange) and nADC (blue) in the motor and visual cortex, as shown in panel (A). Meanwhile, nBOLD (blue) most likely indicates neural deactivation or inhibition in the right angular gyrus, as shown in panel (B). The scale indicates the z-score. The light shaded area on the brain images shows the slab from which data was acquired.

### Exploring Neural Activity/Deactivation with BOLD and ADC maps

Delving deeper into the investigation of whether ADC 0.2/1 fluctuations predominantly reflect neural activity, we analysed the corresponding averaged and normalised timeseries, which were significantly correlated with the task using GLM. Additionally, we compared the ADC brain activation maps with those derived from BOLD imaging. Our objective was to assess whether significant differences could be found in the delays in the timeseries relative to the task and to determine if nADC clusters were localised in similar regions to pBOLD in the activity maps, even when deliberately decoupled from the neurovascular response. Furthermore, we aimed to examine the spatial distribution of nBOLD and pADC clusters to investigate whether they exhibited patterns indicative of neural inhibition or deactivation.

When examining the averaged and normalised significant timeseries derived from ADC and BOLD, it becomes evident that the amplitude of the ADC response is higher than that of the BOLD response. The lower average BOLD amplitude can be attributed to using SE instead of Gradient Echo (GE) EPI. In addition, the higher BOLD sensitivity results in more voxels being identified as significant compared to ADC, thereby reducing the average response amplitude. Furthermore, our results displayed distinct delays between task onset and signal rise/fall, as well as between task offset and return to baseline. Notably, these delays were significantly shorter in nADC compared to pBOLD. This observation resonates with prior research on ADC and comparisons between BOLD and DW MRI timeseries, underscoring DfMRI’s potential sensitivity to neuromorphological coupling [7, 27, 23]. It was proposed that the early onset of the evoked response reflected microstructural changes in tissue [7, 27, 23], considering that it typically takes approximately 2 seconds for blood to traverse from arteries to capillaries and draining veins [57]. Therefore, the ADC and DW MRI timeseries were thought to capture signals prior to the engagement of neurovascular coupling mechanisms. The BOLD signal typically reaches a plateau 6 seconds after stimulus onset and returns to baseline with a similar profile [57], consistent with our finding of a delay of about 5 seconds for the BOLD response. It is important to note that our delay calculation was based on the point where the signal surpasses or falls below its mean, occurring before the signal reaches its plateau, which explains why our observed delay is slightly shorter than the established timeframe.

**Figure 8:**
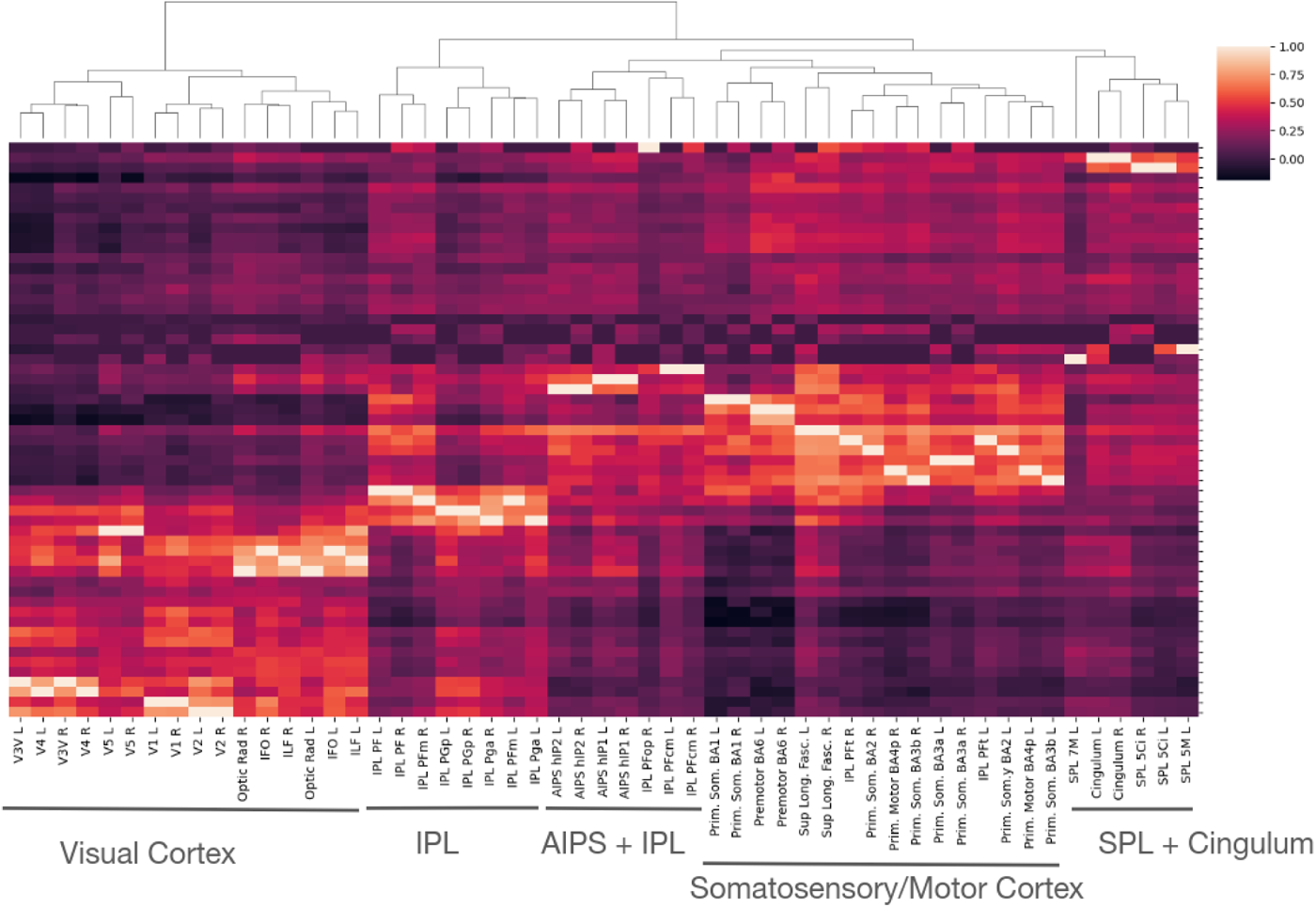
Hierarchical clustering obtained with Pearson partial correlations with the average ADC signal per ROI. The tree shows how the ROIs can be organised based on the similarities of their correlation to the other ROIs (defined by the Juelich and JHU atlases).

Regarding the activity maps, both the averaged subject-level z-score map and the group-level z-score map revealed that pBOLD clusters are located in the visual and motor cortex, as expected. The survival of clusters in the group-level analysis strongly indicates a good alignment of significant clusters across subjects. For the nBOLD, our results indicated that, at the group-level, a significant cluster was located in the right angular gyrus, which is an area of the DMN. This finding provides evidence that nBOLD is sensitive to potential neural deactivation, aligning with previous studies that reported decreased neural activity in DMN areas during goal-directed behaviours [34, 58]. However, likely due to limited sensitivity, nBOLD was only detected in the right angular gyrus. When observing the averaged subject-level analysis, it seemed that the most prominent cluster was situated in the visual cortex but closer to the parietal lobe, in contrast to pBOLD, which was concentrated at the occipital pole. Although this cluster did not emerge in the group-level analysis, its location corresponds to regions of the visual cortex that were identified as non-stimulated during the paradigm and had been previously detected by nBOLD [5].

**Figure 9:**
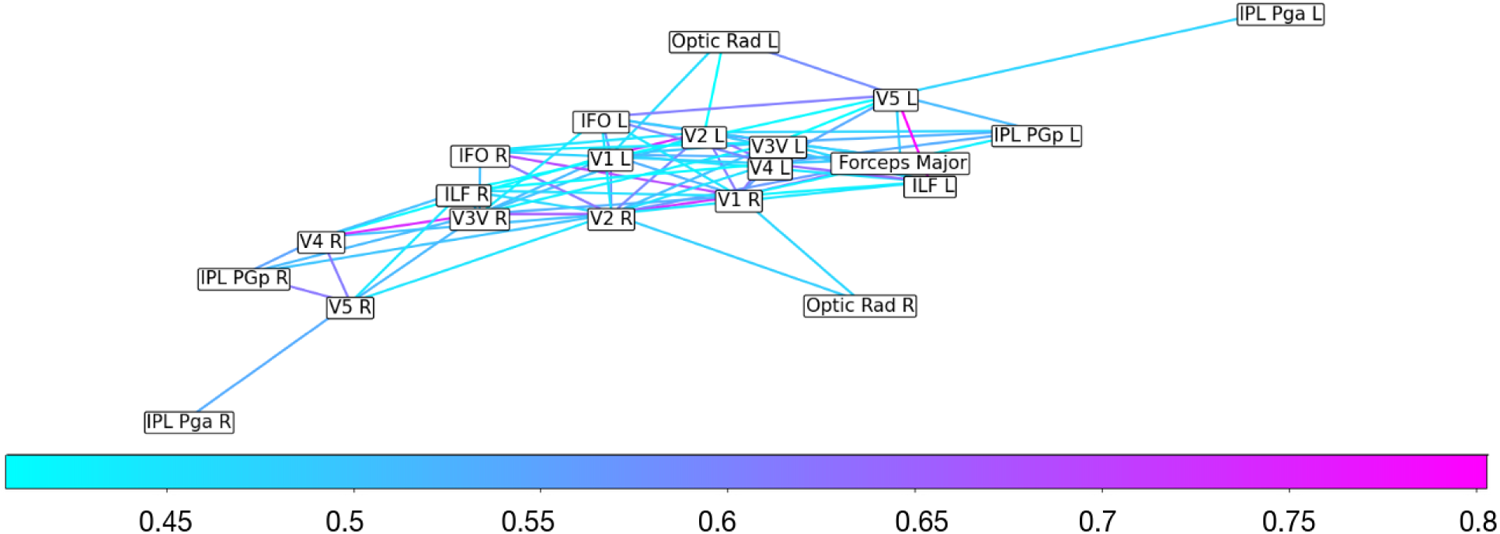
Graph created from the ADC averaged FC matrix. A threshold of 0.4 is applied, so all pairs of ROIs with correlations below this threshold are discarded for the creation of this graph. Only the pairs of ROIs that involved at least one GM ROIs within the visual cortex are shown. The colour scale indicates the partial Pearson correlation value (with the CSF taken as a coviariate).

In the nADC activation maps, clusters in the visual cortex were present in the averaged subject-level analysis but not in the group level analysis. It is likely that the nADC clusters did not survive the group-level analysis in the visual cortex due to the minimal overlap of activated areas between subjects, resulting from low sensitivity and also the small sample size utilised (8 subjects). In the motor cortex, however, a cluster defining a clear path in WM could be distinguished in the group-level analysis, and connected to the motor area depicted by pBOLD clusters. This suggests that nADC could be more sensitive to neural activity in WM tracts than pBOLD, as also previously reported by works examining the ratio of WM-to-GM significant voxels in ADC-fMRI vs BOLD [10]. The slight left-right asymmetry in the motor cortex nADC clusters, particularly evident in the averaged z-score activation map, suggests that ADC-fMRI sensitivity may depend on the angle between the diffusion-encoding direction and the orientations of dendrites and axons. The sensitivity is expected to be maximal to swelling of neuron fibres perpendicular to the diffusion-encoding direction [23]. Furthermore, the imaging slab unfortunately only covers a part of the brain, including the visual and motor cortex. This slab is thus not ideal for studying neural activity within WM tracts such as the CST and the Optic Radiation, that are only partially covered.

In addition to investigating nADC, we explored whether pADC could reveal neural deactivation, serving as a counterpart to nADC, which indicated excitatory patterns. No pADC clusters were significant at the group-level. Although some voxels appeared in the averaged subject-level analysis, they were sparsely distributed throughout the brain. It is likely that pADC signals originate from noise, partial volume artefacts, or potentially from pBOLD contamination, despite our efforts to minimise it.

### Finding evidence of the two-stream hypothesis with FC analysis

Both ADC FC and DW FC were analysed and compared for their effectiveness in organising the ROIs of the brain and in supporting the two-stream hypothesis [59]. According to this hypothesis, information flows from V1 to V2 and then diverges into the ventral stream, directed towards V3v, V4, and the temporal lobe, or into the dorsal stream, directed towards V3d, V3A, V5, and the SPL [60] (see Supplementary Figure 2). Previous investigations into the two-stream hypothesis have employed both structural connectivity [61] and FC with BOLD-fMRI [60]. Therefore, both ADC- and DW-fMRI are promising candidates for such an analysis. It should be noted that our study includes ROIs from both GM and WM, unlike [60]. As a result, an fMRI contrast driven by the vascular response may not be optimal for examining GM-to-WM connectivity. Given the established vascular contributions to the DW timeseries [28], we anticipate that ADC FC will be more effective than DW FC in uncovering insights into GM-to-WM connectivity. Furthermore, ADC-fMRI time courses are less affected by physiological noise (e.g. respiration), which is largely cancelled out when taking the ratio of two consecutive DW images to calculate an ADC. Consequently, we expect ADC FC to yield a better-organised hierarchical clustering of the ROIs even for a low sample size.

In a previous study, the FC strength was shown to decay exponentially with distance indicating that nearby areas connected to one another more strongly than areas farther away [62]. This is in line with the idea that the two pathways are segregated to some extent, and that the nearby areas of the same stream should connect more strongly to one another than to other areas [62]. Unfortunately, the brain slab acquired did not encompass the temporal lobe and only partially covered ILF and IFO WM tracts, posing challenges for studying the ventral stream. Also, the V3d and V3A ROIs were not included in the version of the Juelich atlas used in this study (as opposed to [60]), so a link between V2 and V5 may have been difficult to observe.

The hierarchical clustering derived from the ADC FC provided a remarkably organised depiction of the ROIs within the brain slab, even though the dataset was relatively small (8 subjects). The tree created from this data clearly distinguished between the ROIs of the visual cortex and those of other brain areas. Within this “visual branch”, several sub-groups were evident. One sub-group distinctly separated the V3v/V4 areas from the V5 area, reinforcing the segregation of the ventral and dorsal streams. Another sub-group consisted of the early visual ROIs, V1 and V2, which were involved before the separation into the two streams. The remaining sub-group comprised the WM tracts. These separations highlight the specialised roles these ROIs play in visual processing. The “non-visual branch” also organised the ROIs in a coherent manner. Functionally similar regions were grouped into closely related branches, easily allowing for the identification of several groups based on the ROIs they contained. Additionally, ROIs from the left and right hemispheres tended to be clustered together for most regions, indicating a symmetrical organisation that aligns with the functional similarities between the hemispheres. This structured organisation suggests that ADC-based FC effectively captured functional information, enabling a logical arrangement of ROIs within the hierarchical tree. In contrast, this level of organisation was not observed with DW (b = 0.2 ms/*µm*^2^) based FC (as demonstrated in Supplementary Figure 1) where the ROIs were not arranged in such a coherent manner. This indicates the superior ability of ADC FC to reflect the structural characteristics of brain regions, which is less reliable with DW-based FC. This confirms our initial hypothesis.

Given the high level of organisation demonstrated by the ADC FC of the ROIs, we constructed a FC graph for further investigation based on edges defined by partial Pearson correlations above 0.4 and involving visual ROIs as nodes. The symmetrical organisation within the graph underscored the coherence and integration of the visual processing pathways across both hemispheres. Also, it revealed that the following connections were present in both hemispheres: *V* 1 *→ V* 2 *→ V* 3*v → V* 4 *→ V* 5. Excluding V5, this pattern precisely mirrored the ventral stream. Additionally, the IFO and ILF white matter tracts, known to be involved in the ventral stream [61], were included in this graph and connected to most visual ROIs.

In relation to the dorsal stream, no edges with correlations above 0.4 were found between V5 and the SPL. Instead, we found that V5 was correlated with 2 different ROIs within the IPL, including IPL Pga which is located in the angular gyrus. This observation was consistent with earlier research based on structural MRI, indicating that within the dorsal stream, V5 transmitted information to the angular gyrus in the IPL before it proceeded to the SPL [63, 64, 61].

In summary, despite having obtained data from very few subjects (8 instead of hundreds as in [60]), ADC FC could be used to investigate an organisation amongst visual ROIs and connections that aligned with prior findings on the visual pathways. As expected, ADC FC was more effective than DW FC in organising the ROIs in a logical manner and was also more effective in uncovering information that supports the two-stream hypothesis.

### Limitations

The main limitation of this study was the small sample size, with 8 subjects for ADC and 4 subjects for BOLD. This was particularly challenging for the GLM analysis, as group-level analyses are more robust with larger sample sizes. Additionally, the shortcomings concerning the ADC timeseries in comparison to BOLD primarily stem from their temporal resolution, which are 2 and 1 second(s) respectively. A poorer temporal resolution also poses challenges for the GLM analysis due to the reduced number of time points available and the inability to capture fast processes (high frequencies). The lower temporal resolution could in part be counteracted by using a sliding window of two consecutive images, to calculate an ADC every second. However, this approach would introduce some temporal smoothing and auto-correlations and we have not found it useful in this study. Additionally, given its sensitivity to diffusion along the direction of encoding, ADC may not be optimal for detecting neural activity in voxels containing WM fibres or cortical layers with well-organised dendritic projections that are parallel to this direction. To address this issue, acquiring ADC data using isotropic diffusion encoding could be beneficial. This approach accounts for diffusion in all directions simultaneously, making it independent of the axon fibres’ orientation [65]. Furthermore, it is important to acknowledge the lower Contrast to Noise Ratio (CNR) in ADC timeseries compared to BOLD, which presents challenges in terms of trading off limited sensitivity with higher specificity. Nevertheless, ADC-fMRI identified group-level clusters forming WM pathways that were absent from the BOLD group-level analysis, pointing to potentially higher sensitivity than BOLD in WM. ADC- and BOLD-fMRI could thus be seen as complementary functional contrasts.

Unfortunately, the requirement to maintain a relatively short TR (1 second) for the diffusion protocols necessitated confining our analysis to a brain slab rather than the entire brain. This brain slab was positioned to encompass both the visual and motor cortex, but it did not entirely cover some WM tracts in the visual cortex. With an alternative positioning of the slab, more activity clusters along these WM tracts might have been detected using ADC-fMRI. This limitation could be addressed in the future by increasing the multiband acceleration factor, or employing novel acquisition schemes that allow for higher acceleration factors, thus enabling greater brain coverage within a short TR [66].

## Conclusion and Future works

This study underscores the significance of ADC in the realm of fMRI research, particularly in its capacity to discern areas of morphological alterations indicative of neural swelling during neural activity. ADC effectively addresses inherent limitations of BOLD fMRI, such as the delayed response to task onset and offset, as well as the limited sensitivity to neural activity in WM regions. Notably, when measuring ADC with b-values of 0.2 and 1 ms/*µm*^2^ instead of 0 and 1 ms/*µm*^2^, ADC time courses exhibit reduced susceptibility to contamination from neurovascular coupling.

The presence of significant nADC clusters in regions where neural activity is anticipated, such as the visual and motor cortices, lends weight to the proposition that nADC is intricately intertwined with neural activity. Its sensitivity to WM pathways positions nADC as a potentially excellent complement to pBOLD, which lacks sensitivity in WM regions. Interestingly, pADC clusters were extremely sparse and did not mirror nBOLD ones, suggesting that either nBOLD has a purely vascular origin, or neuronal inhibition cannot be effectively captured by pADC.

At the FC level, ADC-fMRI enabled to not only find evidence of the two-stream hypothesis in the visual system but to organise all the ROIs within the brain slab in a very logical hierarchical tree. This is compelling evidence that ADC-fMRI encompasses essential information on neural activity. The clear tree organisation was possible in spite of the small number of subjects, thanks to the low physiological noise remaining in ADC time courses, which stands in contrast to BOLD or DW time courses, which are more susceptible to physiological noise.

In essence, this comprehensive investigation highlights the potential of ADC-fMRI as a valuable and unique tool in brain functional imaging. It has the capability to provide nuanced insights into neural dynamics and organisational principles within the brain, all while operating independently from neurovascular coupling.

## Acknowledgements

This work was supported by SNSF Spark grant *CRSK −* 2 190882 and ERC Starting Grant ‘FIREPATH’, SERI no.MB22.00032

## References

[1] Kenneth K Kwong, John W Belliveau, David A Chesler, I E Goldberg, Robert M Weisskoff, Bradley P Poncelet, Douglas N Kennedy, Bruce E Hoppel, Michael S Cohen, and Robert Turner. Dynamic magnetic resonance imaging of human brain activity during primary sensory stimulation. Proceedings of the National Academy of Sciences of the United States of America, 89(12):5675–5679, 1992.

[2] S Ogawa, T M Lee, A R Kay, and D W Tank. Brain magnetic resonance imaging with contrast dependent on blood oxygenation. Proceedings of the National Academy of Sciences, 87(24):9868–9872, 1990.

[3] Elizabeth M. C. Hillman. Coupling mechanism and significance of the bold signal: A status report. Annual Review of Neuroscience, 37:161–181, 2014.

[4] R Kawashima, B T O’Sullivan, and P E Roland. Positron-emission tomography studies of cross-modality inhibition in selective attentional tasks: closing the ”mind’s eye”. Proceedings of the National Academy of Sciences, 92(13):5969–5972, 1995.

[5] Ross Wilson, Andrea Thomas, and Stephen D. Mayhew. Spatially congruent negative bold responses to different stimuli do not summate in visual cortex. NeuroImage, 218:116891, 2020.

[6] D. Bressler, N. Spotswood, and D. Whitney. Negative bold fmri response in the visual cortex carries precise stimulus-specific information. PloS one, 2(5):e410, 2007.

[7] Denis Le Bihan, Shin ichi Urayama, Toshihiko Aso, Takashi Hanakawa, and Hidenao Fukuyama. Direct and fast detection of neuronal activation in the human brain with diffusion mri. Proceedings of the National Academy of Sciences, 103(21):8263–8268, 2006.

[8] Jeremy Flint, Brian Hansen, Peter Vestergaard-Poulsen, and Stephen J Blackband. Diffusion weighted magnetic resonance imaging of neuronal activity in the hippocampal slice model. NeuroImage, 46(2):411– 418, 2009.

[9] Tao Jin, Fuqiang Zhao, and Seong-Gi Kim. Sources of functional apparent diffusion coefficient changes investigated by diffusion-weighted spin-echo fmri. Magnetic Resonance in Medicine, 56(6):1283–1292, 2006.

[10] Wiktor Olszowy, Yujian Diao, and Ileana O. Jelescu. Beyond bold: Evidence for diffusion fmri contrast in the human brain distinct from neurovascular response. bioRxiv, 2021.

[11] K^amil Ŭgurbil, Louis Toth, and Dae-Shik Kim. How accurate is magnetic resonance imaging of brain function? Trends in Neurosciences, 26(2):108–114, 2003.

[12] Hirohito Nonaka, Michiaki Akima, Tsutomu Hatori, Toshio Nagayama, Zhen Zhang, and Fujio Ihara. Microvasculature of the human cerebral white matter: arteries of the deep white matter. Neuropathology, 23(2):111–118, 2003.

[13] Dmitry S. Novikov, Jens H. Jensen, Joseph A. Helpern, and Els Fieremans. Revealing mesoscopic structural universality with diffusion. Proceedings of the National Academy of Sciences, 111(14):5088– 5093, 2014.

[14] Leonard B Cohen, Bertil Hille, and Richard D Keynes. Light scattering and birefringence changes during activity in the electric organ of electrophorus electricus. J Physiol, 203(2):489–509, 1969.

15. A. Darquie, J.B. Poline, C. Poupon, H. Saint-Jalmes, and D. Le Bihan. Transient decrease in water diffusion observed in human occipital cortex during visual stimulation. Proc Nat Acad Sci U S A, 98(16):9391–9395, 2002.

[16] Tong Ling, Kevin C. Boyle, Valentina Zuckerman, Thomas Flores, Charu Ramakrishnan, Karl Deisseroth, and Daniel Palanker. High-speed interferometric imaging reveals dynamics of neuronal deformation during the action potential. Proceedings of the National Academy of Sciences, 117(19):10278–10285, 2020.

[17] J Kwon, S Lee, Y Jo, and M Choi. All-optical observation on activity-dependent nanoscale dynamics of myelinated axons. Neurophotonics, 10(1):015003, 2023.

[18] Kuniomi Iwasa, Ichiji Tasaki, and Robert C Gibbons. Swelling of nerve fibers associated with action potentials. Science, 210(4467):338–339, 1980.

[19] Maki Koike-Tani, Takashi Tominaga, Rudolf Oldenbourg, and Tomomi Tani. Instantaneous polarized light imaging reveals activity dependent structural changes of dendrites in mouse hippocampal slices. bioRxiv, 2019.

[20] Erin Walch and Todd A. Fiacco. Honey, i shrunk the extracellular space: Measurements and mechanisms of astrocyte swelling. Glia, 2022.

[21] Thomas Krucker, George R Siggins, and Shelley Halpain. Dynamic actin filaments are required for stable long-term potentiation (ltp) in area ca1 of the hippocampus. Proceedings of the National Academy of Sciences of the United States of America, 97(12):6856–6861, 2000.

[22] Ichiji Tasaki. Rapid structural changes in nerve fibers and cells associated with their excitation processes. The Japanese Journal of Physiology, 49(2):125–138, 1999.

[23] William M Spees, Tsen-Hsuan Lin, and Sheng-Kwei Song. White-matter diffusion fmri of mouse optic nerve. NeuroImage, 65:209–215, 2013.

[24] Tomokazu Tsurugizawa, Luisa Ciobanu, and Denis Le Bihan. Water diffusion in brain cortex closely tracks underlying neuronal activity. Proceedings of the National Academy of Sciences, 110(28):11636– 11641, 2013.

[25] D. Nunes, A. Ianus, and N. Shemesh. Layer-specific connectivity revealed by diffusion-weighted functional mri in the rat thalamocortical pathway. NeuroImage, 184:646–657, 2019.

[26] Essa Yacoub, Kâmil Uludăg, Kâmil Uğurbil, and Noam Harel. Decreases in adc observed in tissue areas during activation in the cat visual cortex at 9.4 t using high diffusion sensitization. Magnetic Resonance Imaging, 26(7):889–896, 2008. Proceedings of the International School on Magnetic Resonance and Brain Function.

[27] Daniel Nunes, Rita Gil, and Noam Shemesh. A rapid-onset diffusion functional mri signal reflects neuromorphological coupling dynamics. NeuroImage, 231:117862, 2021.

[28] Karla L. Miller, Daniel P. Bulte, Hannah Devlin, Matthew D. Robson, Richard G. Wise, Mark W. Woolrich, Peter Jezzard, and Timothy E. J. Behrens. Evidence for a vascular contribution to diffusion fmri at high b value. Proceedings of the National Academy of Sciences, 104(52):20967–20972, 2007.

[29] Daigo Kuroiwa, Takayuki Obata, Hiroshi Kawaguchi, Joonas Autio, Masaya Hirano, Ichio Aoki, Iwao Kanno, and Jeff Kershaw. Signal contributions to heavily diffusion-weighted functional magnetic resonance imaging investigated with multi-se-epi acquisitions. NeuroImage, 98:258–265, 2014.

[30] Joonas A.A. Autio, Jeff Kershaw, Sayaka Shibata, Takayuki Obata, Iwao Kanno, and Ichio Aoki. High b-value diffusion-weighted fmri in a rat forepaw electrostimulation model at 7t. NeuroImage, 57(1):140– 148, 2011.

[31] Andŕe Pampel, Thies H. Jochimsen, and Harald E. Möoller. Bold background gradient contributions in diffusion-weighted fmri—comparison of spin-echo and twice-refocused spin-echo sequences. NMR in Biomedicine, 23(6):610–618, 2010.

[32] R. Nicolas, H. Gros-Dagnac, F. Aubry, and P. Celsis. Comparison of bold, diffusion-weighted fmri and adc-fmri for stimulation of the primary visual system with a block paradigm. Magnetic Resonance Imaging, 39:123–131, 2017.

[33] Alberto De Luca, Lara Schlaffke, Jeroen C. W. Siero, Martijn Froeling, and Alexander Leemans. On the sensitivity of the diffusion mri signal to brain activity in response to a motor cortex paradigm. Human Brain Mapping, 40(17):5069–5082, 2019.

[34] Marcus E Raichle, Ann M MacLeod, Abraham Z Snyder, W James Powers, Debra A Gusnard, and Gordon L Shulman. A default mode of brain function. Proceedings of the National Academy of Sciences, 98(2):676–682, 2001.

[35] Daniel J Felleman and David C Van Essen. Distributed hierarchical processing in the primate cerebral cortex. Cerebral Cortex, 1(1):1–47, 1991.

[36] Robert Desimone, Steven J Schein, James Moran, and Leslie G Ungerleider. Contour, color and shape analysis beyond the striate cortex. Vision Research, 25(3):441–452, 1985.

[37] Mortimer Mishkin, Leslie G. Ungerleider, and Kathleen A. Macko. Object vision and spatial vision: two cortical pathways. Trends in Neurosciences, 6:414–417, 1983.

[38] Leslie G Ungerleider, Mortimer Mishkin, et al. Two cortical visual systems. analysis of visual behavior. *Ingle DJ, Goodale MA*, Mansfield RJW, 1982.

[39] Jesper L Andersson, Stefan Skare, and John Ashburner. How to correct susceptibility distortions in spin-echo echo-planar images: application to diffusion tensor imaging. Neuroimage, 20(2):870–888, 2003.

[40] Jonathan W Peirce. Psychopy–psychophysics software in python. Journal of Neuroscience Methods, 162(1-2):8–13, 2007.

[41] J. Veraart, D. S. Novikov, D. Christiaens, B. Ades-Aron, J. Sijbers, and E. Fieremans. Denoising of diffusion MRI using random matrix theory. Neuroimage, 142:394–406, 2016.

[42] J-Donald Tournier, Robert Smith, David Raffelt, Rami Tabbara, Thijs Dhollander, Maximilian Pietsch, Daan Christiaens, Ben Jeurissen, Chun-Hung Yeh, and Alan Connelly. Mrtrix3: A fast, flexible and open software framework for medical image processing and visualisation. NeuroImage, 202:116137, 2019.

[43] Jessie Mosso, Dunja Simicic, Kadir Simsek, Roland Kreis, Cristina Cudalbu, and Ileana O. Jelescu. Mp-pca denoising for diffusion mrs data: promises and pitfalls. NeuroImage, 263:119634, 2022.

[44] Benjamin Ades-Aron, Gregory Lemberskiy, Jelle Veraart, John Golfinos, Els Fieremans, Dmitry S. Novikov, and Timothy Shepherd. Improved task-based functional mri language mapping in patients with brain tumors through marchenko-pastur principal component analysis denoising. Radiology, 298(2):365– 373, 2021. PMID: 33289611.

[45] Elias Kellner, Bibek Dhital, Valerij G Kiselev, and Marco Reisert. Gibbs-ringing artifact removal based on local subvoxel-shifts: Gibbs-ringing artifact removal. Magnetic Resonance in Medicine, 76:1574–1581, 2016.

[46] Stephen M. Smith, Mark Jenkinson, Mark W. Woolrich, Christian F. Beckmann, Timothy E.J. Behrens, Heidi Johansen-Berg, Peter R. Bannister, Marilena De Luca, Ivana Drobnjak, David E. Flitney, Rami K. Niazy, James Saunders, John Vickers, Yongyue Zhang, Nicola De Stefano, J. Michael Brady, and Paul M. Matthews. Advances in functional and structural mr image analysis and implementation as fsl. NeuroImage, 23:S208–S219, 2004. Mathematics in Brain Imaging.

[47] Brian B Avants, Nicholas J Tustison, Gang Song, Philip A Cook, Arno Klein, and James C Gee. A reproducible evaluation of ants similarity metric performance in brain image registration. Neuroimage, 54(3):2033–2044, 2011.

[48] Bruce Fischl. Freesurfer. Neuroimage, 62 (2):774–781, August 2012. Epub 2012 Jan 10.

[49] Katrin Amunts, Claude Lepage, Laurence Borgeat, Hartmut Mohlberg, Tim Dickscheid, Marie-Eve Rousseau, and Alan C Evans. Bigbrain: An ultrahigh-resolution 3d human brain model. Science, 340(6139):1472–1475, 2013.

[50] Mark W. Woolrich, Brian D. Ripley, Michael Brady, and Stephen M. Smith. Temporal autocorrelation in univariate linear modeling of fmri data. NeuroImage, 14(6):1370–1386, 2001.

[51] Stephen M. Smith, Mark Jenkinson, Mark W. Woolrich, Christian F. Beckmann, Timothy E.J. Behrens, Heidi Johansen-Berg, Peter R. Bannister, Marilena De Luca, Ivana Drobnjak, David E. Flitney, Rami K. Niazy, James Saunders, John Vickers, Yongyue Zhang, Nicola De Stefano, J. Michael Brady, and Paul M. Matthews. Advances in functional and structural mr image analysis and implementation as fsl. NeuroImage, 23:S208–S219, 2004. Mathematics in Brain Imaging.

[52] Mark Jenkinson, Christian F Beckmann, Timothy EJ Behrens, Mark W Woolrich, and Stephen M Smith. Fsl. NeuroImage, 62(2):782–790, 2012.

[53] H. B. Mann and D. R. Whitney. On a test of whether one of two random variables is stochastically larger than the other. The Annals of Mathematical Statistics, 18(1):50–60, 1947.

[54] Mark W. Woolrich, Timothy E.J. Behrens, Christian F. Beckmann, Mark Jenkinson, and Stephen M. Smith. Multilevel linear modelling for fmri group analysis using bayesian inference. NeuroImage, 21(4):1732–1747, 2004.

[55] Susumu Mori, Kenichi Oishi, Hangyi Jiang, Li Jiang, Xin Li, Kazi Akhter, …, and Michael I Miller. Stereotaxic white matter atlas based on diffusion tensor imaging in an icbm template. NeuroImage, 40(2):570–582, 2008.

56. [56] Daniel Müllner. Modern hierarchical, agglomerative clustering algorithms, 2011.

[57] Nikos K. Logothetis. The underpinnings of the BOLD functional magnetic resonance imaging signal. J Neurosci, 23(10):3963–3971, May 2003.

[58] Qolamreza Reza Razlighi. Task-evoked negative bold response in the default mode network does not alter its functional connectivity. Frontiers in Computational Neuroscience, 12:67, 2018.

[59] CT Ferreira, M Ceccaldi, B Giusiano, and M Poncet. Separate visual pathways for perception of actions and objects: evidence from a case of apperceptive agnosia. J Neurol Neurosurg Psychiatry, 65(3):382– 385, 1998.

[60] Byung-Yoon Park, Woo-Young Mark Shim, On James, and Hyeyoung Park. Possible links between the lag structure in visual cortex and visual streams using fmri. Scientific reports, 9(1):4283, 2019.

[61] Sang-Han Choi, Gangwon Jeong, Young-Bo Kim, and Zang-Hee Cho. Proposal for human visual pathway in the extrastriate cortex by fiber tracking method using diffusion-weighted mri. NeuroImage, 220:117145, 2020.

[62] Bhavin R Sheth and Rebecca Young. Two visual pathways in primates based on sampling of space: Exploitation and exploration of visual information. Frontiers in Integrative Neuroscience, 10:37, 2016. Published 2016 Nov 22.

[63] Masahito Uesaki, Hiromasa Takemura, and Hiroshi Ashida. Computational neuroanatomy of human stratum proprium of interparietal sulcus. Brain Structure and Function, 223:489–507, 2018.

[64] Takao Yamasaki and Shozo Tobimatsu. Driving ability in alzheimer disease spectrum: Neural basis, assessment, and potential use of optic flow event-related potentials. Frontiers in Neurology, 9:750, 09 2018.

65. Arthur Spencer, Inès de Riedmatten, Jasmine Nguyen-Duc, Filip Szczepankiewicz, and Ileana Jelescu. Diffusion functional mri with isotropic b-tensor encoding. In ISMRM, Singapore, 2024.

[66] Kawin Setsompop, Qiuyun Fan, Jason Stockmann, Berkin Bilgic, Susie Huang, Stephen F. Cauley, Aapo Nummenmaa, Fuyixue Wang, Yogesh Rathi, Thomas Witzel, and Lawrence L. Wald. High-resolution in vivo diffusion imaging of the human brain with Generalized SLIce Dithered Enhanced Resolution Simultaneous MultiSlice (gSlider-SMS). Magnetic resonance in medicine, 79(1):141, January 2018. Publisher: NIH Public Access.

