## Supplementary Material for "Mapping activity and functional organisation of the motor and visual pathways using ADC-fMRI in the human brain"

| Atlas | Original ROI Name | Short ROI Name |
|---|---|---|
| Juelich | GM Anterior intra-parietal sulcus hIP1 | AIPS hIP1 |
| | GM Anterior intra-parietal sulcus hIP2 | AIPS hIP2 |
| | GM Anterior intra-parietal sulcus hIP3 | AIPS hIP3 |
| | GM Broca's area BA44 | Broca BA44 |
| | GM Broca's area BA45 | Broca BA45 |
| | GM Inferior parietal lobule PF | IPL PF |
| | GM Inferior parietal lobule PFcm | IPL PFcm |
| | GM Inferior parietal lobule PFm | IPL PFm |
| | GM Inferior parietal lobule PFop | IPL PFop |
| | GM Inferior parietal lobule PFt | IPL PFt |
| | GM Inferior parietal lobule Pga | IPL Pga |
| | GM Inferior parietal lobule PGp | IPL PGp |
| | GM Primary auditory cortex TE1.0 | Prim. Aud. TE1.0 |
| | GM Primary auditory cortex TE1.1 | Prim. Aud. TE1.1 |
| | GM Primary auditory cortex TE1.2 | Prim. Aud. TE1.2 |
| | GM Primary motor cortex BA4a | Prim. Motor BA4a |
| | GM Primary motor cortex BA4p | Prim. Motor BA4p |
| | GM Primary somatosensory cortex BA1 | Prim. Som. BA1 |
| | GM Primary somatosensory cortex BA2 | Prim. Som. BA2 |
| | GM Primary somatosensory cortex BA3a | Prim. Som. BA3a |
| | GM Primary somatosensory cortex BA3b | Prim. Som. BA3b |
| | GM Secondary somatosensory cortex / Parietal operculum OP1 | Sec. Som. OP1 |
| | GM Secondary somatosensory cortex / Parietal operculum OP2 | Sec. Som. OP2 |
| | GM Secondary somatosensory cortex / Parietal operculum OP3 | Sec. Som. OP3 |
| | GM Secondary somatosensory cortex / Parietal operculum OP4 | Sec. Som. OP4 |
| | GM Superior parietal lobule 5Ci | SPL 5Ci |
| | GM Superior parietal lobule 5L | SPL 5L |
| | GM Superior parietal lobule 5M | SPL 5M |
| | GM Superior parietal lobule 7A | SPL 7A |
| | GM Superior parietal lobule 7M | SPL 7M |
| | GM Superior parietal lobule 7PC | SPL 7PC |
| | GM Superior parietal lobule 7P | SPL 7P |
| | GM Visual cortex V1 BA17 | V1 |
| | GM Visual cortex V2 BA18 | V2 |
| | GM Visual cortex V3V | V3V |
| | GM Visual cortex V4 | V4 |
| | GM Visual cortex V5 | V5 |
| | GM Premotor cortex BA6 | Premotor BA6 |
| | WM Optic radiation | Optic Rad |
| | GM Insula Ig1 | Insula Ig1 |
| JHU | Corticospinal tract | Corticospinal |
| | Cingulum (cingulate gyrus) | Cingulum |
| | Cingulum (hippocampus) | Cing. Hipp. |
| | Forceps major | Forceps Major |
| | Inferior fronto-occipital fasciculus | IFO |
| | Inferior longitudinal fasciculus | ILF |
| | Superior longitudinal fasciculus | Sup Long. Fasc. |

Table S1: Names of ROIs defines by the Juelich and JHU that were utilised in this work.

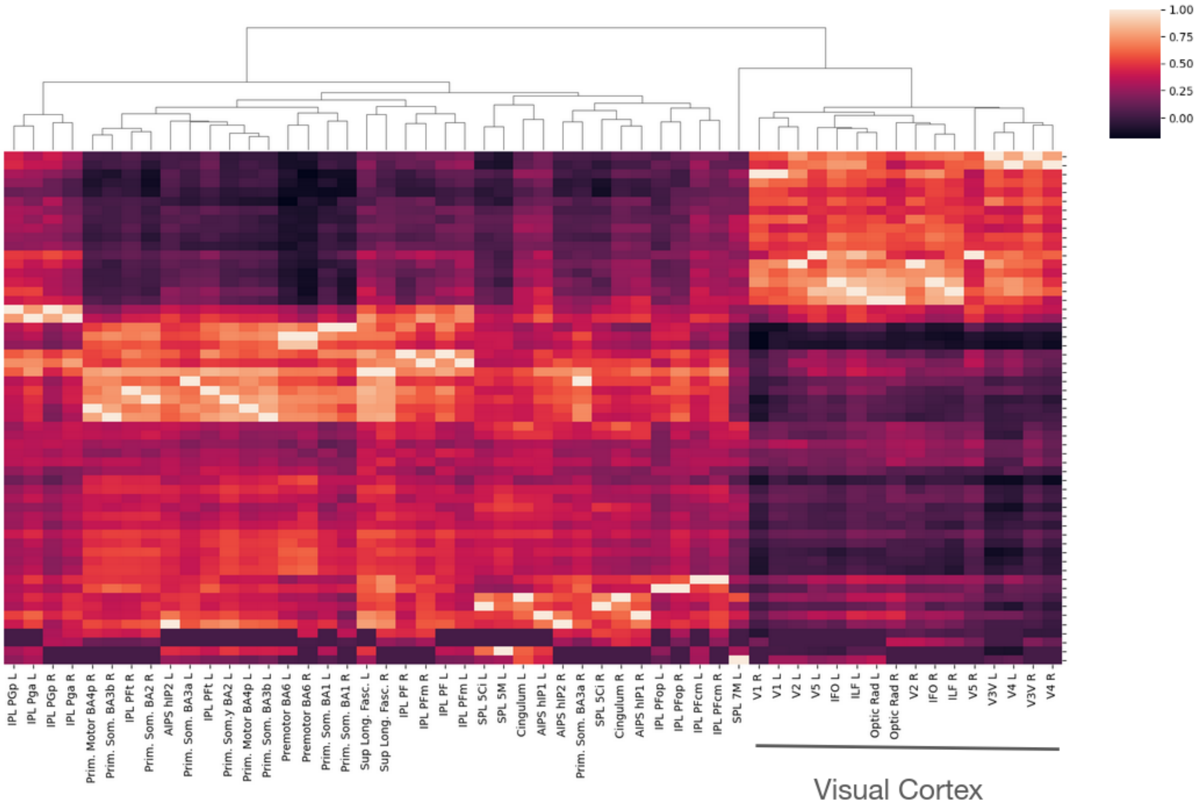


Figure S1: Hierarchical clustering obtained from the FC matrix with Pearson partial correlations with the average DW (b = 0.2 ms/$\mu m^2$) signal per ROI. The tree shows how the ROIs can be organised based on the similarities of their correlation to the other ROIs (defined by the Juelich and JHU atlases)

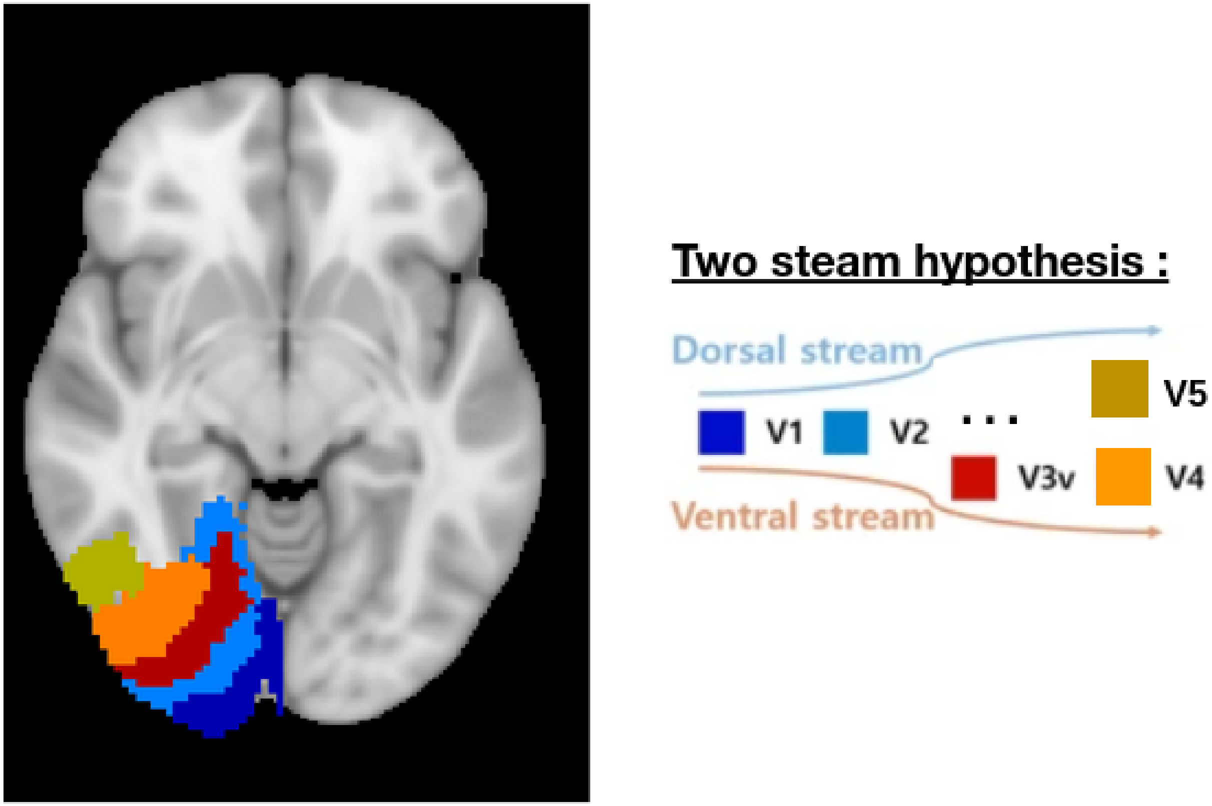


Figure S2: Illustration of V1, V2, V3v, V4 and V5 areas in one hemisphere, as defined by the Juelich atlas. A scheme, modified from Park et al. [59], illustrates the two-stream hypothesis.
